# Nanocarrier-driven dual targeting VCAM-1/Collagen IV enables RNA interference-mediated silencing of Smad3 and Runx2 to mitigate aortic valve disease

**DOI:** 10.1101/2025.07.31.668019

**Authors:** Geanina Voicu, Cristina Ana Mocanu, Florentina Safciuc, Maria Anghelache, Mihaela Turtoi, Mariana Deleanu, Maya Simionescu, Ileana Manduteanu, Manuela Calin

**Author notes:** Correspondence: Manuela Calin “Medical and Pharmaceutical Bionanotechnologies” Laboratory, Institute of Cellular Biology and Pathology “Nicolae Simionescu” of the Romanian Academy, 050568 Bucharest, Romania.

## Abstract

**BACKGROUND:** Calcific aortic valve disease (CAVD) is a common malady with few treatment options other than valve replacement by surgery or transcatheter aortic valve implantation (TAVI). Endothelial-to-mesenchymal transition (EndMT) of valvular endothelial cells and osteogenic differentiation of valvular interstitial cells are crucial processes of CAVD. Smad3 and Runx2 are key transcription factors (TFs) that drive these processes by regulating gene expression and cellular functions. We hypothesize that downregulation of these TFs with nanoparticle-mediated RNA interference could mitigate aortic valve stenosis and calcification.

**METHODS:** We engineered dual-targeted lipid-polymer hybrid nanocarriers (lipopolyplexes, LPP) to deliver short-hairpin RNA (shRNA) for gene silencing in pathologically remodeled aortic valve. The nanocarriers simultaneously target vascular cell adhesion molecule-1 (VCAM-1) and collagen IV, enhancing specificity toward inflamed and fibrotic valvular tissue. Encapsulated shRNA constructs were designed to silence either Smad3 or Runx2 (yielding formulations V/Cp-LPP/shSmad3 and V/Cp-LPP/shRunx2). Therapeutic efficacy was evaluated in a mouse model of atherosclerosis aggravated by diabetes, mimicking the pathological environment of CAVD.

**RESULTS:** The dual-targeted lipopolyplexes effectively facilitated gene delivery to the aortic valve, ensuring efficient transfection. Treatment with V/Cp-LPP/shSmad3 and V/Cp-LPP/shRunx2 resulted in marked silencing of Smad3 and Runx2, accompanied by significant suppression of osteogenic markers, including osteopontin, alkaline phosphatase, and osteocalcin, as well as reduced αSMA expression in valve tissue. Our data further identify Runx2 as a novel upstream modulator of Smad3 expression, unveiling a previously unrecognized Runx2-Smad3 regulatory axis with important implications for valvular pathology and targeted therapy. Beyond localized effects, systemic administration of these lipopolyplexes led to reduced plasma concentrations of alkaline phosphatase, cholesterol, and triglycerides, while maintaining hepatic and renal function, suggesting additional benefits on systemic metabolic homeostasis.

**CONCLUSIONS:** These findings highlight the pivotal role of Smad3 and Runx2 downregulation in mitigating aortic valve calcification, unveiling both molecules as compelling therapeutic targets in CAVD.

**Highlights:**

- Lipopolyplexes dual-targeted to VCAM-1 and Collagen IV successfully transfect valvular cells and drive plasmid expression in the valve leaflets of a CAVD mouse model.
- Therapeutic delivery of shRNA specific for Smad3 and Runx2 led to significant downregulation of osteogenic markers (osteopontin, alkaline phosphatase, osteocalcin) and reduced αSMA in the aortic valve.
- The study establishes Runx2 as an upstream regulator of Smad3 expression with implications for valve disease intervention.
- The engineered lipopolyplexes demonstrated a favorable safety profile and systemic benefits: decreased plasma levels of alkaline phosphatase, cholesterol, and triglycerides.
- The study validates the lipopolyplexes-based strategy as a safe and targeted platform for localized molecular reprogramming in calcific aortic valve disease.

## Introduction

Calcific aortic valve disease (CAVD) is becoming a common malady around the world, leading to significant health issues and a high risk of death.^1^ It is characterized by hemodynamic perturbation of the blood flow to the aorta due to the fibrosis and calcification of valve leaflets.^2^ In the early stage of CAVD, activation and localized disruption of the valvular endothelial cells (VEC) layer allow subendothelial recruitment of immune cells via upregulated cell adhesion molecules, such as intercellular adhesion molecule (ICAM)-1, vascular cell adhesion molecule (VCAM)-1, and E-selectin.^3^ VEC also undergoes an endothelial-to-mesenchymal transition (EndMT) and acquires mesenchymal markers, such as α-Smooth Muscle Actin (αSMA), Snai1, and Octamer-binding transcription factor (OCT)4.^4^ An important transcription factor (TF) involved in inflammation and EndMT processes is the suppressor of mothers against decapentaplegic (Smad) 3, which regulates the transcriptional activity of Zinc finger E-box binding homeobox (Zeb) 1/2, Twist1, and Snai1/2, and favors the acquisition of a mesenchymal phenotype.^5^ Proinflammatory cytokines released by EndMT-engaged VEC and immune cells induce the osteodifferentiation of valvular interstitial cells (VIC) and the increased activity of Runt-related transcription factor (Runx) 2, which causes the overexpression of osteogenic proteins such as osteopontin (OSP), alkaline phosphatase (ALP), osteocalcin (OSC), and bone morphogenic proteins (BMP) in these cells.^6^ Notably, both hypercholesterolemia and diabetes are well-established risk factors for CAVD. Elevated expression of Runx2, OSP, and ALP has been observed in the aortic valves of rabbits with hypercholesterolemia-induced atherosclerosis^7^, as well as in patients with CAVD and comorbid diabetes mellitus^8^.

Despite advances in surgical and transcatheter valve replacements, the mortality risk of CAVD increases every five years.^9^ Additionally, the therapies currently in use do not effectively slow down the progression of the disease. Thus, identifying promising therapies that target key processes involved in CAVD initiation and progression is essential.^10^ As a strategy to reduce aortic valve calcification, we planned to develop an efficient delivery system for short hairpin (sh)RNA targeting two proteins that are overexpressed in the dysfunctional aortic valve, VCAM-1 and collagen IV. Our approach utilizes nanoparticles to silence either Smad3 or Runx2, effectively preventing the endothelial-to-mesenchymal transition (EndMT), the osteodifferentiation of VIC, or both. In previous studies, we demonstrated that the progression of CAVD is featured by endothelial activation and increased VCAM-1 expression,^11^ and, also, by a disturbance in the synthesis of extracellular matrix proteins (collagen IV)^12^; with time, the latter become exposed to the blood circulation after endothelial denudation.^13,14^ Thus, VCAM-1 and collagen IV may serve as effective targets for drug delivery systems to treat CAVD.

Lipopolyplexes are lipid/polymer-based, biocompatible, and stable colloidal systems,^15^ well-suited for *in vivo* administration.^12^ They are relatively easy to synthesize, with precise control over size and surface charge. It has been reported that lipopolyplexes have numerous advantages as delivery systems for RNA interference (RNAi), such as colloidal stability, reduced cytotoxicity, and high transfection efficiency.^16^ Lipopolyplexes carrying shRNA plasmids, specific for Runx2 and Smad3, complexed with fullerene (C60)-polyethyleneimine (PEI), mitigate the osteodifferentiation of VIC^12^ and reverse the EndMT process of VEC^11^ *in vitro*. Other studies showed that Runx2 ubiquitination via Forkhead box protein O (FOXO) 1, followed by degradation,^17^ deacetylation, and nuclear export of Runx2 mediated by Sirtuin (SIRT) 6^18^ and Runx2 silencing by direct binding of Twist1 to its promotor^19^ inhibit human aortic valve calcification. Long noncoding RNA lncTSI^20^ and BBT-877, an inhibitor of autotaxin (lysophospholipase D),^21^ reduce Smad3 phosphorylation and Runx2 level, thus preventing the progression of CAVD. Therefore, our goal was to downregulate either Smad3 or Runx2 expression by delivering specific shRNA encapsulated in lipopolyplexes to the aortic valve of a mouse model for CAVD. To enhance the delivery of lipopolyplexes to the dysfunctional aortic valve, their surface was decorated with two peptides recognizing VCAM-1 and collagen IV (V/Cp-LPP/shSmad3, V/Cp-LPP/shRunx2). Employing the CAVD mouse model, the experiments showed that, upon i.v. injection, V/Cp-LPP/shSmad3 and V/Cp-LPP/shRunx2 lipopolyplexes suppress the expression of Smad3 and Runx2 in the aortic valve, along with their downstream effector molecules: ALP, OSP, and OSC. This molecular silencing was accompanied by reduced lipid deposition, lower ALP enzymatic activity within the valve leaflets, and decreased circulating ALP levels. Taken together, these findings underscore the therapeutic promise of selectively silencing Smad3 and Runx2, and position targeted gene knockdown as a compelling strategy to halt the progression of CAVD.

## Materials and methods

### Reagents

Detailed information on key reagents is provided in the Major Resources Table (**Table S4**), in the Supplemental Materials.

### Animal model

Recently, we introduced the diabetic ApoE-deficient mice fed a high-fat diet as a suitable experimental murine model for studying early molecular and functional perturbations in aortic valves.^22^ On this model of diabetic/atherosclerotic mice, we demonstrated that the dysfunctional aortic valve is characterized by an increased level of VCAM-1^11^ and the enhanced deposition of collagen IV in the extracellular matrix.^12^ Now, we employed this murine model to search for the therapeutic effect of double-targeted lipopolyplexes carrying shRNA plasmids specific for Runx2 or Smad3. Twelve-week-old ApoE-deficient (B6.129P2-*Apoe^tm1Unc^* N11) and C57BL/6 (C57BL/6NTac) mice from Taconic Biosciences were used for the experiments. The animals had access to a standard rodent diet and water *ad libitum* and were housed in individually ventilated cages in a specific pathogen-free facility (SPF) on a 12-hour light/dark cycle. To induce diabetes and accelerate the progression of aortic valve calcification, the animals were injected intraperitoneally with 55 mg/kg streptozotocin (STZ) in a total volume of 150 µL 0.9% NaCl for 5 consecutive days. The final dose of STZ was administered concurrently with the initiation of a high-fat diet, comprising standard chow enriched with 1% cholesterol and 15% butter (82% fat),^23^ maintained for up to one or three weeks. The weight of the animals and the glycemia were monitored as welfare parameters.

The study protocols and experiments were approved by the Ethics Committee of the Institute of Cellular Biology and Pathology “Nicolae Simionescu” and by the National Sanitary Veterinary and Food Safety Authority, authorization no. 448/April 02, 2019. The research was conducted following the Romanian Law no. 43/2014 (Official Monitor, Part I no. 326, pages 2– 4), which transposes the EU Directive 2010/63/EU on the protection of animals used for scientific purposes. The study follows the principles outlined in the ARRIVE guidelines, ensuring transparency and reproducibility in reporting. ^24^ Group sizes were determined based on a 30% anticipated difference in mean between groups with α = 0.05, a statistical power of 90%, using the free power calculator provided by the Laboratory Animal Services Centre (LASEC). The standard deviation used in sample size calculation was obtained from previous experiments. The analysis suggested an optimum sample size of n = 8 mice per group (with a minimum of n = 6). However, the experimental groups were established at n = 9 each, after considering the adverse effects induced by STZ administration.

### Preparation of VCAM-1/ Collagen IV double-targeted lipopolyplexes encapsulating shRNA plasmids

#### shRNA sequences

To downregulate the mRNA expression of Runx2 and Smad3, a mix of five MISSION®shRNA Plasmid DNA shRNA sequences targeting the murine Runx2 gene (cat. no. SHCLND-NM_009820.2) and Smad3 gene (cat. no. SHCLND-NM_016769.3), respectively, was used. The specific shRNA sequences for the murine Runx2 and Smad3 genes from the RNAi Consortium (TRC) Version 1 library are presented in **Table S1**. As a control plasmid, MISSION® pLKO.1-puro non-Mammalian shRNA Control Plasmid DNA (shCtr) was employed. *Escherichia coli* host strain DH5α was used for plasmid amplification; the isolation and purification were performed using the GenElute-Plasmid Midiprep kit (Sigma-Aldrich, Germany).

#### Lipopolyplexes synthesis

The lipopolyplexes (LPP) were prepared using the reverse-phase evaporation method following previously established protocols.^15^ To prepare C60-PEI/shRNA polyplexes at an N/P ratio of 25 (ratio of nitrogen atoms in C60-PEI to phosphorus atoms in DNA^25^), conjugated fullerene (C60)-polyethyleneimine (PEI) and shRNA plasmids were each diluted in equal volumes of 2 × HB buffer (20 mM HEPES, 10% D-glucose, pH = 7.4). The solutions were then adjusted to a final volume of 15 mL with HB buffer (10 mM Hepes, 5% D-glucose, pH = 7.4). After a 10-minute incubation at room temperature, the mixture was briefly vortexed to form cationic polyplexes with an N/P ratio of 25. Preformed cationic polyplexes were mixed with 3 mM anionic DOPG diluted in chloroform/methanol solution (ratio 2:1, v/v). After 30 minutes at room temperature, 15 mL of chloroform and distilled water (1:1, v/v) were added to the reverse micelles and centrifuged at 830 × g for 7 minutes. The aqueous phase was removed, and 6.7 mM POPC, 0.1 mM Mal-PEG-DSPE, 0.1 mM Metoxy-PEG-DSPE (dissolved in chloroform), and 15 mL of HB buffer were added to the organic phase. The lipid-coated polyplexes (lipopolyplexes) were then vortexed vigorously and sonicated for one minute. The chloroform was evaporated under a vacuum using a rotary evaporator (Laborota 4000, Heidolph, Schwabach, Germany), at 37°C, in a round-bottom glass bottle. A hand extruder and polycarbonate membranes of 200 nm and 100 nm from Avanti Polar Lipids (Alabaster, AL, USA) were used for aqueous dispersion extrusion and for obtaining lipopolyplexes uniform in size (LPP/shRunx2, LPP/shSmad3, and LPP/shCtr).

#### Surface decoration of lipopolyplexes with peptides recognizing VCAM-1 and Collagen IV

To enhance the targeting capability, double-targeted nanoparticles were developed by surface functionalization with two peptides, one selective for VCAM-1 (V, sequence NH_2_-VHPKQHRGGSKGC-COOH) and the other for collagen IV (Cp, sequence NH2-KLWVLPKGGGC-COOH), combined at a 1:1 ratio. The procedure for peptide coupling is as follows. First, the disulfide bonds in peptides were broken by adding a reducing agent (TCEP) for 2 hours at room temperature. The excess TCEP was removed by dialysis overnight at 4°C against coupling buffer (10 mM Na_2_HPO_4_, 10 mM NaH_2_PO_4_, 2 mM EDTA, 30 mM NaCl, pH 6.7) using a dialysis membrane of 100-500 Da. Then, the peptides were added to the lipopolyplexes and incubated overnight at 4°C to form the bonds between the maleimide group at the distal end of lipid Mal-PEG-DSPE contained in the liposomes’ bilayer and cysteines from the carboxy tail of the peptides. After saturation of the uncoupled maleimide groups with 1 mM L-cysteine for 30 minutes, the lipopolyplexes were further centrifuged using Amicon centrifugal filter units of 100 kDa to separate the free peptides from peptides-coupled lipopolyplexes.

The concentration of peptides specific for VCAM-1 and collagen IV coupled on the surface of lipopolyplexes was indirectly determined by quantifying the uncoupled peptides using Ultra-High-Performance-Liquid-Chromatography (UHPLC), as described elsewhere.^26^ The encapsulation efficiency of plasmids in the LPP was detected using the Quant-iT™ PicoGreen® dsDNA kit (cat. no. R11490, ThermoFischer Scientific), as previously reported.^15^

#### Characterization of lipopolyplexes

The hydrodynamic diameter and Zeta potential of lipopolyplexes were determined as previously mentioned.^15^ The physical stability of lipopolyplexes stored at 4°C for 3 weeks was checked at fixed time intervals (1, 2, and 3 weeks) by measuring the hydrodynamic diameter and the Zeta potential. The results were analyzed using the built-in Zetasizer Software 7.12 (Malvern Instruments, Malvern, UK).

### Transfection efficiency of double-targeted lipopolyplexes in murine aortic valves

To evaluate the specificity for dysfunctional aortic valve and the transfection efficiency of VCAM-1/ Collagen IV double-targeted lipopolyplexes as delivery vectors for shRNA plasmids, diabetic ApoE-deficient mice (n = 2) and C57BL/6 mice (n = 2) were i.v. injected with V/Cp-LPP/pEYFP lipopolyplexes containing 10 µmols lipids/1.5 mg pEYFP per kg body weight (100 μL lipopolyplexes per mouse). The pEYFP plasmid is derived from *Aequorea Victoria* and encodes a green/yellow fluorescent protein. After 48 hours, the aortic roots were harvested, cryoprotected, and sliced for microscopy investigation under a fluorescence microscope. The nuclei were stained with ProLong Gold Antifade Mountant with DAPI. The fluorescence images were acquired using an Olympus IX81 inverted microscope (Shinjuku City, Tokyo, Japan) at 20× and 40× magnification, and image processing was performed using Fiji Is Just ImageJ (NIH), a free software program version 1.8.0.

### Assessment of the level of osteogenic molecules after lipopolyplexes administration in the mouse model of CAVD

Diabetic ApoE-deficient mice placed on a high-fat diet (HFD) for three weeks were divided into four experimental groups, with 9 mice per group, totaling 36 mice. The mice were i.v. injected two times a week for the last two weeks (4 doses) with VCAM-1/ Collagen IV double-targeted lipopolyplexes carrying a mix of five shRNA plasmid sequences targeting mouse Smad3 or Runx2 mRNA (10 µmols lipids/1.5 mg shRNA plasmid per kg body weight), in a volume of 100 μL V/Cp-LPP/shSmad3 or V/Cp-LPP/shRunx2 per mouse. As controls, the administration of double-targeted lipopolyplexes containing shCtr plasmids (V/Cp-LPP/shCtr) and PBS was employed. At 48 hours after the last i.v. injection, the animals were anesthetized, exsanguinated via open heart puncture, and perfused with PBS. The plasma separated after blood centrifugation was stored at −80°C until further assays. The aortic valve, lungs, spleen, liver and kidneys were collected and processed for mRNA expression determinations, cryoprotection and histological staining.

#### Quantitative Real-Time Polymerase Chain Reaction (qRT-PCR)

The mRNA expression of genes involved in EndMT (Smad3) and calcification (Runx2, ALP, OSP, OSC) of the murine aortic valve was determined by quantitative real-time polymerase chain reaction (qRT-PCR). The total mRNA was extracted in TRIzol™ reagent from tissue homogenate of aortic valves (n = 6 aortic valves/experimental group, divided into 3 pools of 2 valves). Separately, the mRNA expression of Smad3 and Runx2 genes was also determined in pulmonary, hepatic, and renal tissue homogenates (20-40 mg of tissue/probe). The tissue homogenate was obtained using a UPH200H probe-type sonicator from Hielscher (Hielscher Ultrasonics GmbH, Teltow, Germany). The concentration of the isolated RNA was determined using a Spectrophotometer NanoDrop 1000 (ThermoFisher Scientific, Waltham, MA, USA). First-strand cDNA synthesis was performed employing 2 μg of total RNA and MMLV reverse transcriptase, according to the manufacturer’s protocol (Invitrogen, Waltham, Massachusetts, USA). Gene expression quantification was performed using a Viia7 RT-PCR System (ThermoFisher Scientific Inc., Waltham, MA, USA), SYBR Green I chemistry, and primers for mouse genes reported in **Table S2**. The mRNA fold changes were quantified using the 2^−ΔΔCT^ method, normalized to β-actin (ACTB), and expressed relative to levels found in the control mice group (injected with PBS), considered as 1.

To determine whether the downregulation of Runx2 in VIC, the primary cells involved in valve calcification, directly affects Smad3 gene expression, we conducted qRT-PCR on cells activated in a medium containing 25 mM glucose and osteogenic factors (50 µg/mL ascorbic acid, 10 mM β-glycerophosphate, 10 nM dexamethasone) (HGOM). The experimental protocol for the *in vitro* analysis is presented in the **Supplemental material**.

#### Immunofluorescence

The protein expression of molecules involved in CAVD was evaluated by immunofluorescent staining of murine aortic valve cryosections. Thus, the slides with valve cryosections obtained as described elsewhere^12^ were rehydrated with warm PBS and incubated overnight at 4°C with the following primary antibodies: rabbit polyclonal antibody anti-Smad2/3 (1:100, cat. no. ab217553), rabbit polyclonal antibody anti-Runx2 (1:100, cat. no. PA1-41519), rabbit polyclonal antibody anti-osteopontin (1:100, cat. no. PA5-34579), goat polyclonal antibody anti-alkaline phosphatase (1:40, cat. no. AF2910), rabbit polyclonal antibody anti-periostin (1:1000, cat. no. ab14041) diluted in 3% BSA/PBS. Before incubation with the anti-Runx2 antibody, samples underwent heat-induced epitope retrieval (HIER) to unmask antigenic sites. This was achieved by treating slides with 50 mM Tris-base buffer (pH 9.0) at 96 °C for 3 hours.^23^ The next day, the valve sections were incubated with the appropriate secondary antibody: Alexa Fluor 594 donkey anti-rabbit (cat. no. A21207), Alexa Fluor 594 donkey anti-goat (cat. no. A11058), diluted 1:1000 in 3% BSA/PBS for 1 hour at room temperature. In addition we used a double-staining protocol to visualize the colocalization of CD31 and αSMA. For this protocol, the slides were incubated ON at 4°C with the first primary antibody, rabbit anti-CD31 (cat. no. PA5-29166, ThermoFisher Scientific, 1:500 dilution), followed by incubation with the secondary antibody, anti-rabbit coupled with Alexa Fluor 594, for one hour at RT. The incubation with the second primary antibody, namely goat anti-αSMA (cat. no. PA5-18292, ThermoFisher Scientific, 1:100 dilution), was also performed ON, and the secondary antibody used was FITC-conjugated rabbit anti-goat antibody (cat. no. A16143, Invitrogen, 1:1000 dilution). After washing with PBS, the sections were mounted in ProLong Gold Antifade Mountant with DAPI. Stained sections were visualized under a fluorescence microscope (Olympus IX81 equipped with an XM10 camera and a TRITC filter). For each sample, a total of 2-3 images were taken at 40× magnification. The image processing was performed using ImageJ (NIH, USA) freeware program version 1.8.0.

#### Alkaline phosphatase activity in the aortic valve

Cryosections of aortic valves were stained for alkaline phosphatase activity using the method described elsewhere.^27^ Slides containing aortic valve sections were rinsed for 30 minutes with warm PBS, then incubated for 1 hour at 37°C in a solution of 2% sodium 5,5-diethyl barbiturate, 3% β-sodium glycerophosphate, 2.7% CaCl₂, 5% MgSO₄ × 7H₂O, and distilled water, adjusted to pH 9.4. Next, the sections were washed three times with alkaline water and incubated with 2% cobalt nitrate for 5 minutes. After rinsing with distilled water, the slides were immersed in 0.5% ammonium sulfide for 20 minutes, then thoroughly washed with distilled water, 100% ethanol, and xylene before being mounted in Canada balsam for preservation. The images were taken using a camera-equipped Olympus IX81 inverted microscope at 20× magnification. Two slices per mouse were morphometrically quantified for measuring the alkaline phosphatase activity area using Fiji Is Just ImageJ (NIH) freeware program version 1.8.0. The area positive for alkaline phosphatase activity was calculated as a percentage of the total surface area of the aortic valve.

### Ultrasound-based cardiac function analysis in the experimental model of CAVD treated with lipopolyplexes

The aortic valve function of diabetic and hyperlipidemic ApoE-deficient mice injected i.v. with lipopolyplexes (V/Cp-LPP/shSmad3, V/Cp-LPP/shRunx2 and V/Cp-LPP/shCtr) or PBS was evaluated using a high-resolution ultrasonic imaging system for small animals (Vevo2100, FUJIFILM VisualSonics, Toronto, ON, Canada). After shaving the mice’s chest, they were placed under light anesthesia with 2% isoflurane and maintained on a heated platform, with continuous monitoring of heart rate and core temperature. Pulsed wave-Doppler (PW-Doppler) mode was utilized to measure transvalvular velocity across the aortic valve and the left ventricular outflow tract velocity time integral (LVOT VTI) using parasternal long-axis views. VTI and velocity serve as key indicators of cardiac output and systolic function. The mean gradient was determined by averaging instantaneous gradients over the ejection period, derived from transaortic velocity. Additionally, cardiac output and stroke volume were assessed using both M- and PW-Doppler modes. To establish potential changes in heart rate and valve structure, we conducted preliminary echocardiography after diabetes induction and a final scan just before sacrifice. Measurements were extracted from recorded videos and automatically calculated using specific formulas in VevoLab300 software.

### Quantification of plasma osteogenic factors in a mouse model of CAVD treated with lipopolyplexes

Blood was collected in EDTA tubes (5 mM) and the plasma was obtained by centrifugation at 2000 rpm for 10 minutes at 4°C, then stored at −80°C until the assays were performed. The osteogenic molecules ALP (cat. no. AF2910, R&D Systems), OSC, and OSP (cat. no. RK03088, RK03094, ABclonal Biotechnology) were quantified in 1:10 diluted plasma using an ELISA assay, adhering to the manufacturer’s guidelines. The optical density was detected using an Infinite® M200 PRO microplate reader spectrophotometer (Tecan, Männedorf, Switzerland) at the specified wavelength for optimal detection. The concentrations of ALP, OSC, and OSP were determined by plotting their respective standard curves and performing calculations based on the generated data.

### Safety evaluation of lipopolyplexes

#### Biochemical plasma parameters

The biochemical measurements of plasma parameters were performed using DiaLab reagent kits for cholesterol (cat. no. D95116), triglycerides (cat. no. D00389), plasma urea (BUN; cat. no. 402999), creatinine (cat. no. D95595), transaminases aspartate aminotransferase (AST; cat. no. D94610), alanine aminotransferase (ALT; cat. no. D94620), according to manufacturer’s instructions. After pipetting the probes into a 96-well plate, the appropriate volumes of reagents were added and the absorbance at 578 nm was measured using an Infinite® M200 PRO microplate reader spectrophotometer (Tecan, Männedorf, Switzerland). The parameter concentrations were determined through simultaneous measurements using the recommended standards, Assayed Universal Calibration Serum (cat. no. D98485, Diacal Autor), ensuring accuracy and consistency.

#### Histopathological analysis of murine pulmonary, hepatic, renal, and splenic tissues

The pulmonary, hepatic, renal, and splenic tissues were fixed in 4% PFA/PBS overnight and subsequently cryoprotected following the previously described protocol. The 5 µm cryosections (three sections per slide) were washed with PBS and stained with hematoxylin and eosin (cat. no. H-3502, VECTOR Laboratories), following the manufacturer’s instructions for optimal staining quality. The micrographs were taken using an Olympus IX81 inverted microscope equipped with a camera. In pulmonary sections, the alveolar septal wall thickness and the number of nuclei, as a measure of cellular density, were quantified by analyzing optical microscopy micrographs taken with a 20× objective using ImageJ 1.53c freeware (NIH, USA) according to the macros mentioned previously.^23^ Ten random fields were used per mouse.

### Statistical analysis

The results are expressed as mean ± standard deviation (S.D.). Statistical analyses were performed using GraphPad Prism software v. 10.4.1 (GraphPad Software Inc., San Diego, CA, USA). Statistical differences were evaluated using one-way ANOVA with the Tukey post hoc test. For echocardiographic measurements and *in vitro* RT-PCR analysis, Sidak’s multiple comparisons test was employed. Statistically significance of differences: *p < 0.05, ** p < 0.01, *** p < 0.001.

## Results

### Characterization of lipopolyplexes carrying shRNA-plasmids

The average hydrodynamic diameter immediately after synthesis of V/Cp-LPP/shRunx2 and V/Cp-LPP/shCtr was ∼ 140 nm, the size of V/Cp-LPP/shSmad3 was around 200 nm, while that of untargeted Scr-LPP/shCtr was ∼ 230 nm (**Figure 1A**). The polydispersity index (PDI) below 0.25 indicates that lipopolyplexes are homogenous and unimodally distributed. The Zeta potential measurements showed that all lipopolyplexes are stable negatively charged nanoparticles, with values between −23 ÷ −33 mV. Over a storage period of up to three weeks at 4°C, the lipopolyplexes exhibited minimal changes in hydrodynamic diameter and Zeta potential, demonstrating their relative stability as a nanoparticle dispersion.

**Figure 1.**
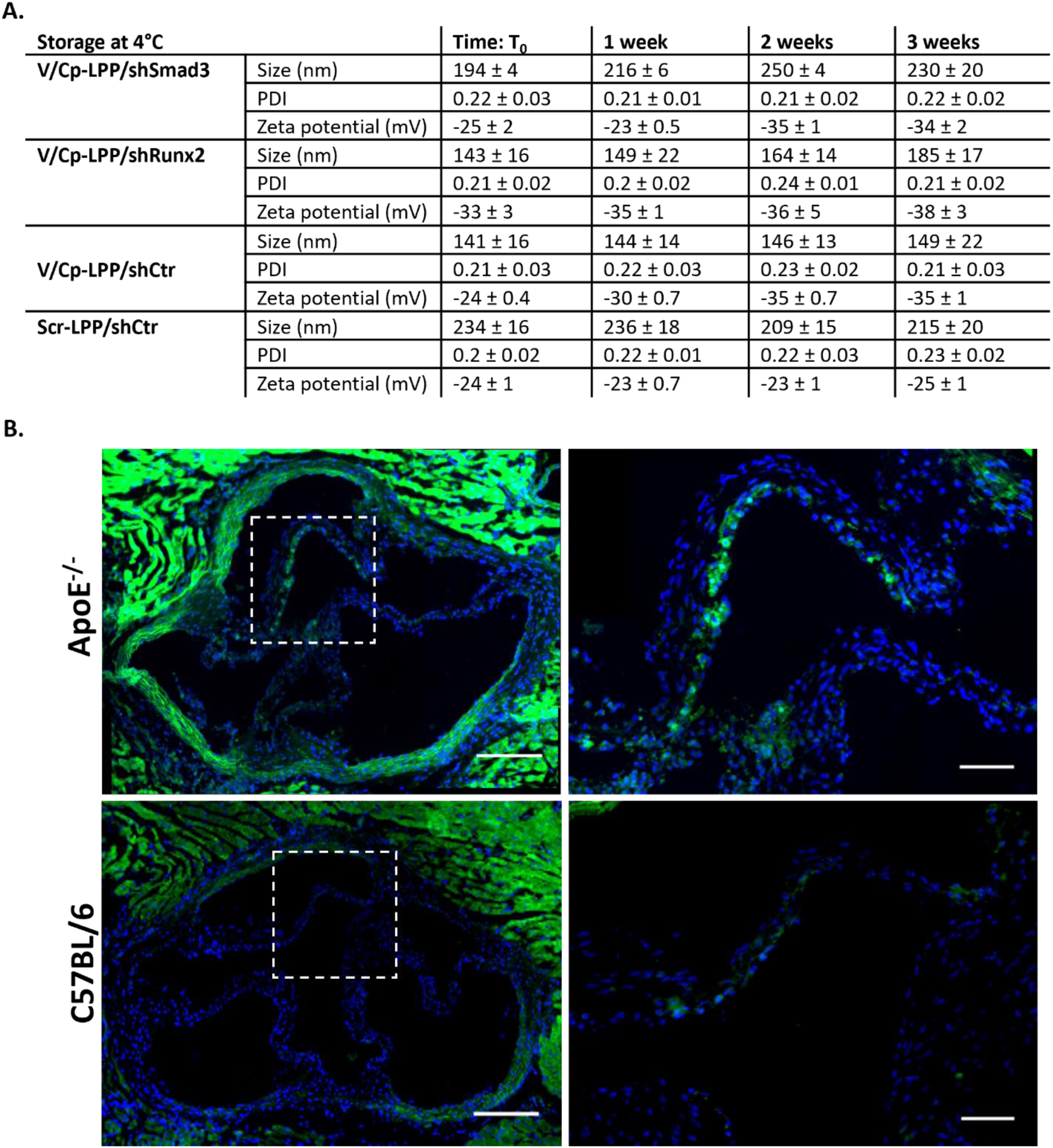
**A.** Characterization of lipopolyplexes. Hydrodynamic diameter, Zeta potential, and polydispersity index of V/Cp-LPP/shSmad3, V/Cp-LPP/shRunx2, V/Cp-LPP/shCtr, and Scr-LPP/shCtr. Results are reported as mean ± S.D. from three records representing the average of 13 internal measurements. **B.** Representative images of YFP fluorescence in the aortic valve sections of diabetic/hyperlipidemic ApoE-deficient and C57BL/6 mice after i.v. injection of V/Cp-LPP/pEYFP lipopolyplexes; the whole valve is shown with a scale bar of 200 µm. The region illustrated for high magnification is marked with a square; the detail is shown with a scale bar of 100 µm. Nuclei are labeled with DAPI (blue). The image processing was performed using Fiji Is Just ImageJ (NIH) freeware program version 1.8.0.

The HPLC data measurements indicated an amount of ∼ 7.2 µg VCAM-1 peptide and ∼ 8 µg collagen IV peptide per µmol lipid coupled to the surface of lipopolyplexes. The amount of encapsulated fluorescently labeled shRNA plasmid quantified using Quant-iT PicoGreen reagent was ∼ 7.55 µg Smad3 shRNA plasmid (75.5% encapsulation efficiency), ∼ 9.79 µg Runx2 shRNA plasmid (98% encapsulation efficiency), and ∼ 7.6 µg shCtr plasmid (76% encapsulation efficiency) encapsulated per µmol lipid.

### Double-targeted lipopolyplexes efficiently transfect the aortic valve in a CAVD mouse model

To assess the delivery and transfection efficiency of double-targeted lipopolyplexes in the dysfunctional aortic valve, diabetic/hyperlipidemic ApoE-deficient and wild-type C57BL/6 mice received retro-orbital injections of V/Cp-LPP/pEYFP lipopolyplexes, enabling targeted evaluation of their ability to deliver the pEYFP plasmid encoding yellow fluorescent protein (YFP). Forty-eight hours after injection, the aortic valve leaflets of diabetic/hyperlipidemic ApoE-deficient mice exhibited stronger YFP expression than those from C57BL/6 mice **(Figure 1B**). The findings demonstrated that VCAM-1/ Collagen IV double-targeted lipopolyplexes could serve as highly efficient carriers for the *in vivo* delivery of plasmids to the dysfunctional aortic valve, ensuring targeted gene transfer and potential therapeutic benefits. Their stability and transfection capability highlight their suitability for precise genetic modulation within the affected valve tissue.

### Lipopolyplexes-mediated silencing of Smad3 and Runx2 reduces the expression of osteogenic molecules in the aortic valve of the CAVD mouse model

We next investigated the therapeutic effects of lipopolyplexes carrying the shRNA-Smad3 and shRNA-Runx2 plasmids following administration in mice, assessing their potential to reduce osteogenic molecule expression and impede the progression of aortic valve calcification. To induce dysfunction of the aortic valve in ApoE-deficient mice, a combination of intraperitoneal streptozotocin injections for five days and a high-fat diet for three weeks was applied. Subsequently, the mice received retro-orbital injections with different types of lipopolyplexes two times per week for two weeks, while PBS-injected mice served as controls. Forty-eight hours after the last injection, the therapeutic effect of lipopolyplexes was investigated in the aortic valve by measuring Smad3, Runx2, ALP, OSC, and OSP gene expression. A significant reduction in Smad3 gene expression, approximately 3.5-fold, was determined following treatment with V/Cp-LPP/shSmad3 or V/Cp-LPP/shRunx2 in mice compared to control groups, as illustrated in **Figure 2A**. The RT-PCR results were supported by fluorescence microscopy images (**Figure 2B**), where a substantial reduction of ∼ 18-fold was assessed in animal groups treated with targeted lipopolyplexes containing shSmad3 or shRunx2 compared to groups that received PBS or V/Cp-LPP/shCtr (**Figure 2C**). Runx2 gene expression was reduced in mice receiving V/Cp-LPP/shRunx2 by ∼ 1.7-fold compared to control groups (**Figure 2A**). The most significant reduction in mRNA-Runx2 expression, approximately a tenfold decrease compared to the control, was observed when V/Cp-LPP/shSmad3 was utilized, demonstrating its superior efficacy in gene silencing of Runx2. The gene-level findings translated into corresponding protein expression for lipopolyplexes containing the shRunx2 plasmid (**Figure 2B**); however, this was not observed for those containing the shSmad3 plasmid, where the reduction in Runx2 protein levels did not align with the trend of mRNA expression.

**Figure 2.**
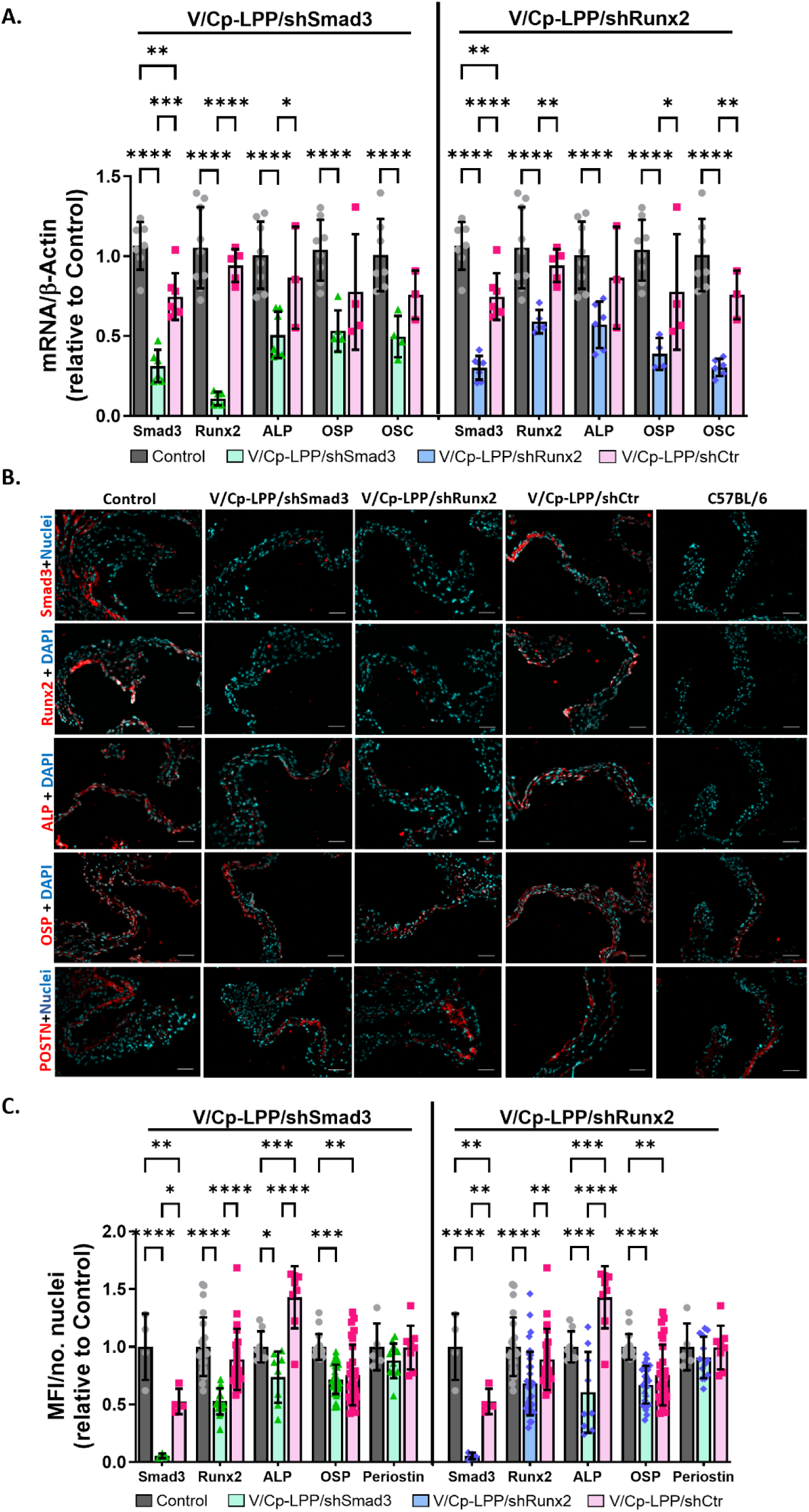
Expression of osteogenic genes and their corresponding proteins in the aortic valve of lipopolyplexes-treated mice. **A.** mRNA expression of Smad3, Runx2, alkaline phosphatase (ALP), osteopontin (OSP), and osteocalcin (OSC) in the aortic valve of diabetic/hyperlipidemic ApoE-deficient mice injected with VCAM-1/ Collagen IV double-targeted lipopolyplexes encapsulating shRNA-Smad3, shRNA-Runx2, or control plasmids (notations: V/Cp-LPP/shSmad3, V/Cp-LPP/shRunx2, V/Cp-LPP/shCtr). Mice receiving PBS were used as controls. The results, normalized to β-actin and relative to the control group (considered as 1), are represented as mean ± S.D. For each of the biological replicates (n = 3 pooled samples from 2 valves), 2-3 technical replicates were performed. **B.** Representative fluorescence images indicating the presence of investigated proteins: Smad3, Runx2, ALP, OSP, and periostin (POSTN) (red) in the aortic valve. Nuclei are stained with DAPI (cyan). Scale bar: 50 µm. **C.** Smad3, Runx2, ALP, OSP, and POSTN protein level quantification on fluorescence microscopy images. The results are represented as Mean Fluorescence Intensity (MFI), normalized to the number of nuclei and relative to the control group. Two to five fields per sample (n = 3 mice/experimental group) were analyzed. Statistical significances (one-way ANOVA with the Tukey post hoc test): * p < 0.05, ** p < 0.01, *** p < 0.001, **** p < 0.0001. The image processing was performed using Fiji Is Just ImageJ (NIH) freeware program version 1.8.0.

To dissect the *in vivo* results and further investigate the co-dependence between the transcription factors Smad3 and Runx2, we aimed to analyze the effects of lipopolyplexes on a cell population actively involved in aortic valve stenosis and calcification, namely valvular interstitial cells (VIC). Thus, we conducted qRT-PCR on VIC incubated in an activatory medium and treated with collagen IV-targeted lipopolyplexes carrying shRunx2 plasmid (Cp-LPP/shRunx2). Briefly, the cells cultured for 5 days in medium containing a normal concentration of glucose, 5 mM (NG) or osteogenic medium (HGOM) containing 25 mM glucose and osteogenic factors (50 μg/mL ascorbic acid, 10 mM β-glycerophosphate, 10 nM dexamethasone) were treated for 48 hours with 1 µg of shRNA-Runx2 (sequence for human Runx2: GCTACCTATCACAGAGCAATT) plasmid encapsulated in Cp-LPP/shRunx2. The results, shown in **Figure S1A**, indicate that HGOM-osteodifferentiated VIC exhibit an approximately 50% and 30% increase in mRNA-Runx2 and mRNA-Smad3 levels, respectively, compared to control, non-activated VIC. However, the incubation for 48 hours of osteodifferentiated cells with Cp-LPP/shRunx2 reduced the levels of both Runx2 and Smad3 mRNA by ∼50% and ∼70%, respectively, compared to HGOM-exposed cells without lipopolyplexes treatment. This result aligns with *in vivo* studies and suggests that the downregulation of Runx2 directly affects Smad3 expression in VIC, indicating a regulatory link between the two genes.

Furthermore, we evaluated the aortic valves from groups of mice treated with lipopolyplexes for the mRNA level of additional osteogenic molecules, namely ALP, OSC, and OSP, whose expression is regulated by Smad3 and Runx2 and plays a crucial role in valve calcification (**Figure 2A**). The treatment with V/Cp-LPP/shSmad3 led to an approximate two-fold downregulation in the mRNA level of ALP, OSP, and OSC. Besides, following treatment with V/Cp-LPP/shRunx2, we observed an approximate 1.8-fold reduction in ALP mRNA levels in mice. Additionally, the mRNA levels of OSP and OSC were significantly decreased by approximately 2.5-fold and 3.5-fold, respectively, compared to control mice. V/Cp-LPP/shCtr treatment did not alter gene expression of any investigated molecule in the aortic valve compared to the control mice.

Protein expression of osteogenic molecules was assessed in cryosections of murine aortic valve tissue using fluorescence microscopy (**Figure 2B**). Downregulation of Smad3 level in the aortic valves of mice injected with V/Cp-LPP/shSmad3 resulted in a ∼1.35-fold reduction in ALP protein expression compared with the control group (**Figure 2C**). Additionally, OSP levels in these mice decreased by 1.4-fold relative to controls. In the aortic valves of mice treated with V/Cp-LPP/shRunx2, ALP and OSP protein expression levels were reduced by approximately 1.6-fold and 1.5-fold, respectively, confirming the modulatory role of Runx2 in the transcription of osteogenic molecules. POSTN protein expression remained unchanged following treatment with V/Cp-LPP/shSmad3 or V/Cp-LPP/shRunx2. Additionally, administration of V/Cp-LPP/shCtr did not further alter the levels of osteogenic molecules compared to the control group.

### ALP activity in the aortic valve of the mouse model with CAVD following V/Cp-LPP/shSmad3 and V/Cp-LPP/shRunx2 treatments

The aortic valve of the mouse model of CAVD was analyzed in all experimental groups following i.v. administration of V/Cp-LPP/shSmad3, V/Cp-LPP/shRunx2, V/Cp-LPP/shCtr lipopolyplexes, and PBS. ALP activity in valve cryosections was assessed by staining with ammonium sulfide, which yields a black cobalt sulfide precipitate in regions of enzymatic activity. Enhanced ALP activity was observed across the cross-sectional area of aortic valve samples from both control mice (PBS-injected) and those treated with V/Cp-LPP/shCtr, as indicated by extensive regions of multilayered black cobalt sulfide precipitate (**Figure 3A**). In mice treated with V/Cp-LPP/shSmad3 and V/Cp-LPP/shRunx2, ALP activity in the aortic valves was reduced by approximately 30%, though the change did not reach statistical significance (**Figure 3B**). This reduction was visually corroborated by a diminished accumulation of black cobalt sulfide precipitate in the valve tissue (**Figure 3A**).

**Figure 3.**
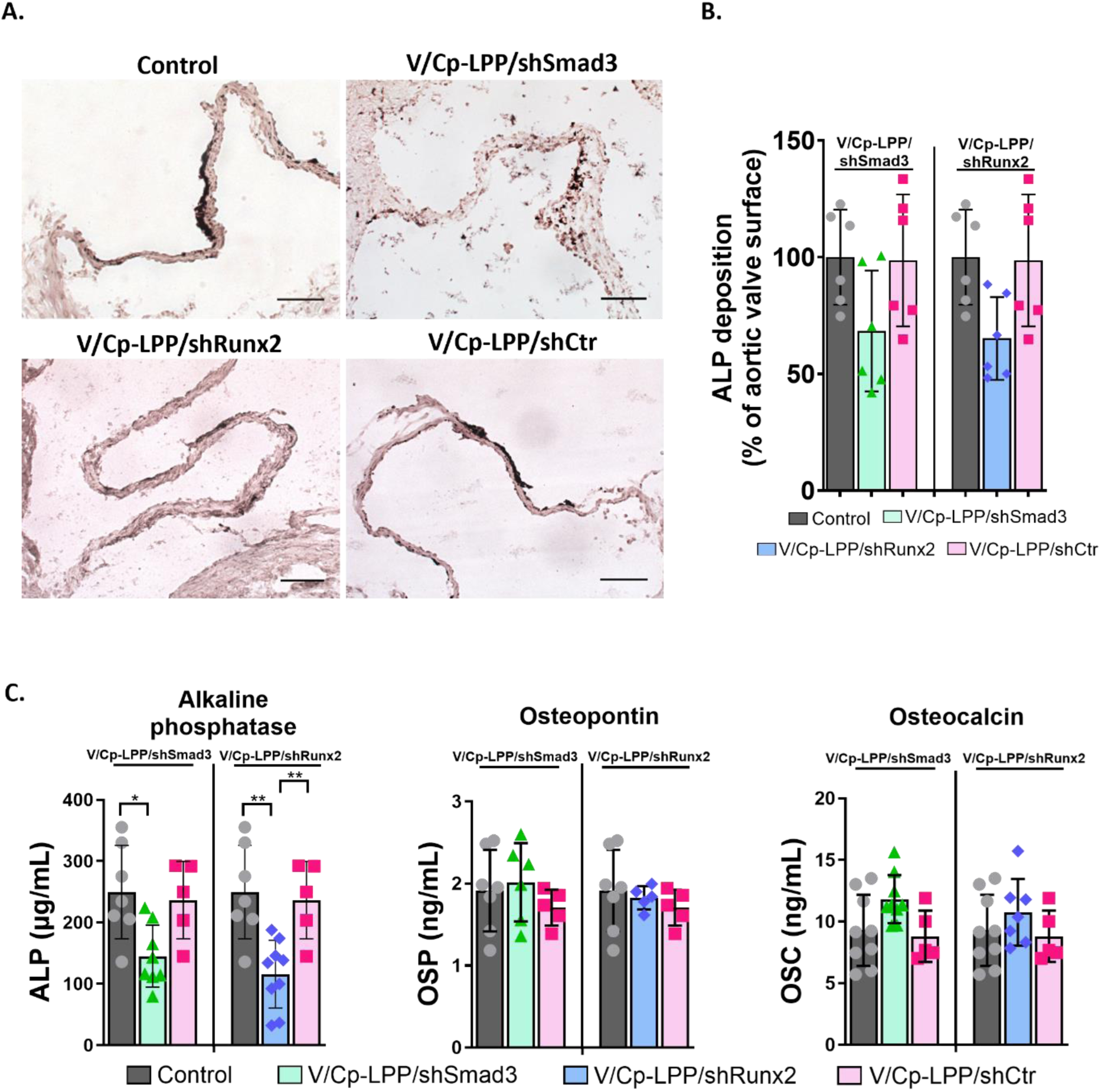
**A.** Representative images of alkaline phosphatase activity (black deposits) in the aortic valve of diabetic/hyperlipidemic ApoE-deficient mice injected with VCAM-1/ Collagen IV double-targeted lipopolyplexes encapsulating shRNA-Smad3, shRNA-Runx2, or control plasmids (notations: V/Cp-LPP/shSmad3, V/Cp-LPP/shRunx2, V/Cp-LPP/shCtr). Mice receiving PBS were used as controls. Scale bar: 500 µm. **B.** Quantification of alkaline phosphatase activity area calculated as a percent of total aortic valve surface area. Results are the means ± S.D. of two sections per sample (n = 3 mice/experimental group). The image processing was performed using Fiji Is Just ImageJ (NIH) freeware program version 1.8.0. **C.** Levels of plasmatic osteogenic molecules (ALP, OSP, OSC) measured in mice injected with V/Cp-LPP/shSmad3, V/Cp-LPP/shRunx2, V/Cp-LPP/shCtr lipopolyplexes, and control mice. Data are mean ± S.D. (n = 5-9 samples per experimental group). Statistical significances (one-way ANOVA with the Tukey post hoc test): * p < 0.05, ** p < 0.01.

### Assessment of osteogenic molecules in the plasma of CAVD mice

To further define the therapeutic efficiency of downregulating transcriptional factors Smad3 and Runx2, we examined the plasma osteogenic molecules ALP, OSP and OSC levels (**Figure 3C**). Mice injected with V/Cp-LPP/shSmad3 and V/Cp-LPP/shRunx2 had statistically lower levels of ALP detected in their plasma, by ∼ 1.7-fold (* p < 0.05) and 2.2-fold (** p < 0.01), respectively, compared to control mice. No changes in the ALP level were detected when V/Cp-LPP/shCtr lipopolyplexes were injected. The levels of OSP and OSC did not vary statistically in any of the mouse experimental groups compared to control mice injected with PBS.

### Investigation of the EndMT process in the aortic valve of the mouse model of CAVD

Since Smad3 plays a crucial role in EndMT, a process in which endothelial cells acquire mesenchymal characteristics, contributing to fibrosis and aortic valve dysfunction, investigating how treatment with V/Cp-LPP/shSmad3 influences this pathway is essential for understanding its potential therapeutic effects in mitigating pathological tissue remodeling. Our findings in cultured VIC demonstrate that Runx2 plays a regulatory role in the expression of Smad3 (**Figure S1A**), revealing a crucial link between these transcription factors. Building on this insight, we further explored the effects of V/Cp-LPP/shRunx2 treatment on EndMT in the aortic valve to clarify its impact on the Runx2–Smad3 axis and assess its therapeutic relevance. We analyzed CD31 and αSMA as distinct markers of endothelial and mesenchymal cells, respectively, to evaluate the effect of treatment with targeted lipopolyplexes on their expression and potential role in modulating cellular transitions in the aortic valve of the mouse model of CAVD.

Merged fluorescent images revealed colocalization of CD31 with αSMA in the aortic valves of the CAVD mouse model, indicating that EndMT is actively triggered in this pathological setting (**Figure 4A**). In the group of mice treated with V/Cp-LPP/shSmad3, CD31 protein expression increased by approximately 40%, while αSMA protein expression decreased by 15%; however, these changes did not reach statistical significance (**Figure 4B)**. While Smad3 silencing via V/Cp-LPP/shSmad3 did not produce statistically significant changes, the trend toward elevated CD31 and diminished αSMA expression suggests a potential modulatory influence on EndMT.

**Figure 4.**
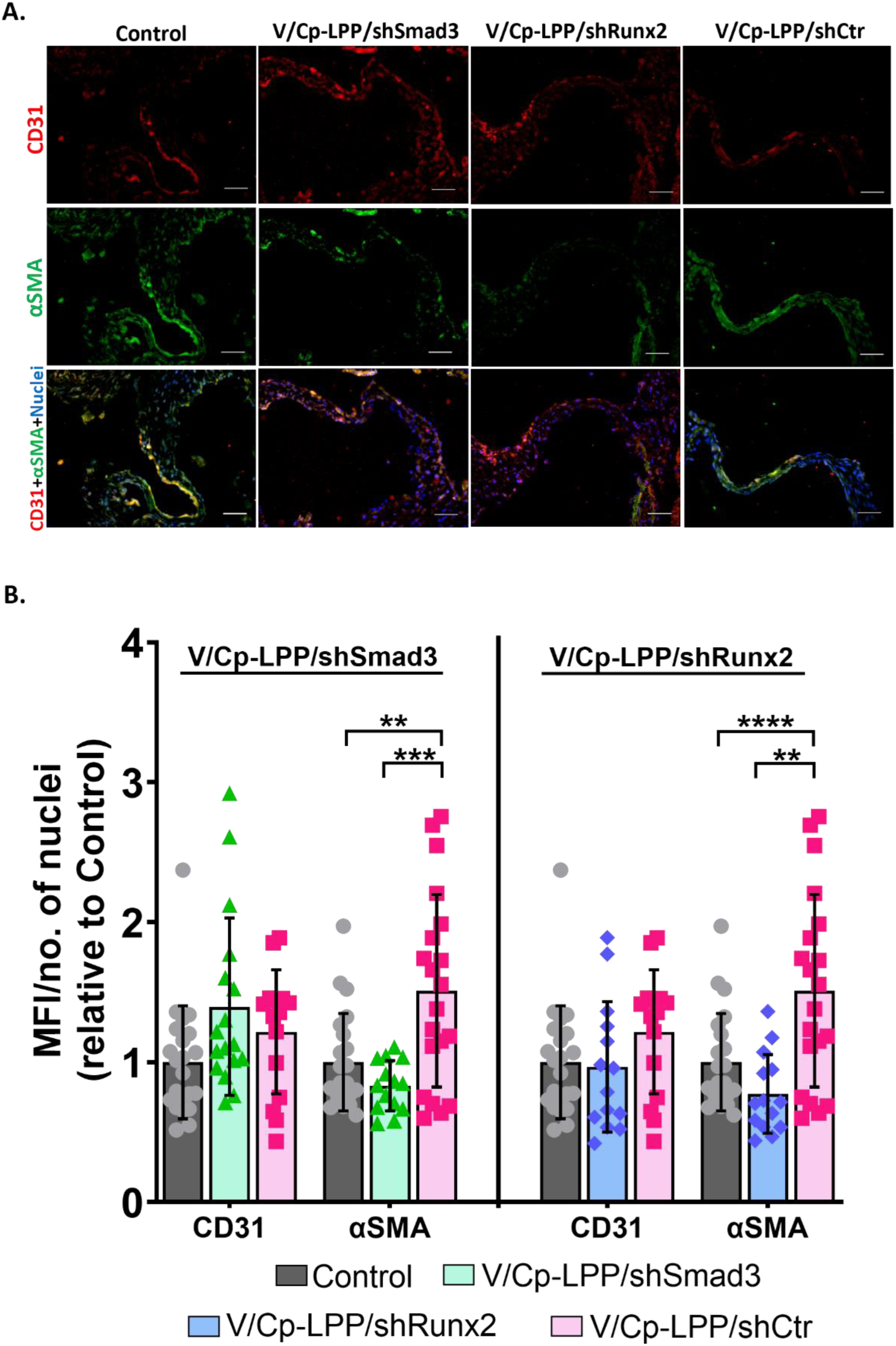
**A.** Representative fluorescence images showing the colocalization of CD31 (red) and αSMA (green) in the aortic valves of diabetic/hyperlipidemic ApoE-deficient mice injected with V/Cp-LPP/shSmad3, V/Cp-LPP/shRunx2, and V/Cp-LPP/shCtr. Nuclei are stained with DAPI (blue). Scale bar: 50 µm. **B.** Quantification of CD31 and αSMA protein expression on fluorescence microscopy images. The results are represented as Mean Fluorescence Intensity (MFI), normalized to the number of nuclei and relative to the control group. Five to seven fields per sample (n = 3 mice/experimental group) were analyzed. Statistical significances (one-way ANOVA with the Tukey post hoc test): * p < 0.05, ** p < 0.01, *** p < 0.001, **** p < 0.0001. The image processing was performed using Fiji Is Just ImageJ (NIH) freeware program version 1.8.0.

Although the treatment did not significantly alter CD31 levels, the trend toward reduced αSMA expression (by 20%) following V/Cp-LPP/shRunx2 treatment suggests that Runx2 may play a regulatory role in mesenchymal marker expression, potentially impacting EndMT dynamics. The absence of a concomitant increase in endothelial markers may suggest that the EndMT process may be only partially modulated, rather than fully reversed. This likely reflects an early or incomplete transition blockade, potentially due to the short duration of induction.

Despite that CD31 expression remained unchanged following treatment with shCtr plasmid encapsulated in lipopolyplexes, an unexpected increase in αSMA was observed. This effect may stem from sequence-independent effects of the control plasmid itself. Non-targeting shRNAs and plasmid DNA can modulate gene expression through off-target mechanisms, including microRNA and transcriptional interference.^28^

### Echocardiographic assessment in the mouse model of CAVD

**Figure 5A** illustrates the changes in blood glucose concentrations measured at multiple time points in ApoE-deficient mice: before STZ administration (T0), after STZ administration (Ti), and before sacrifice (Tf). Mice administered STZ were classified as diabetic when the blood glucose levels consistently exceeded 250 mg/dL throughout the study.

**Figure 5.**
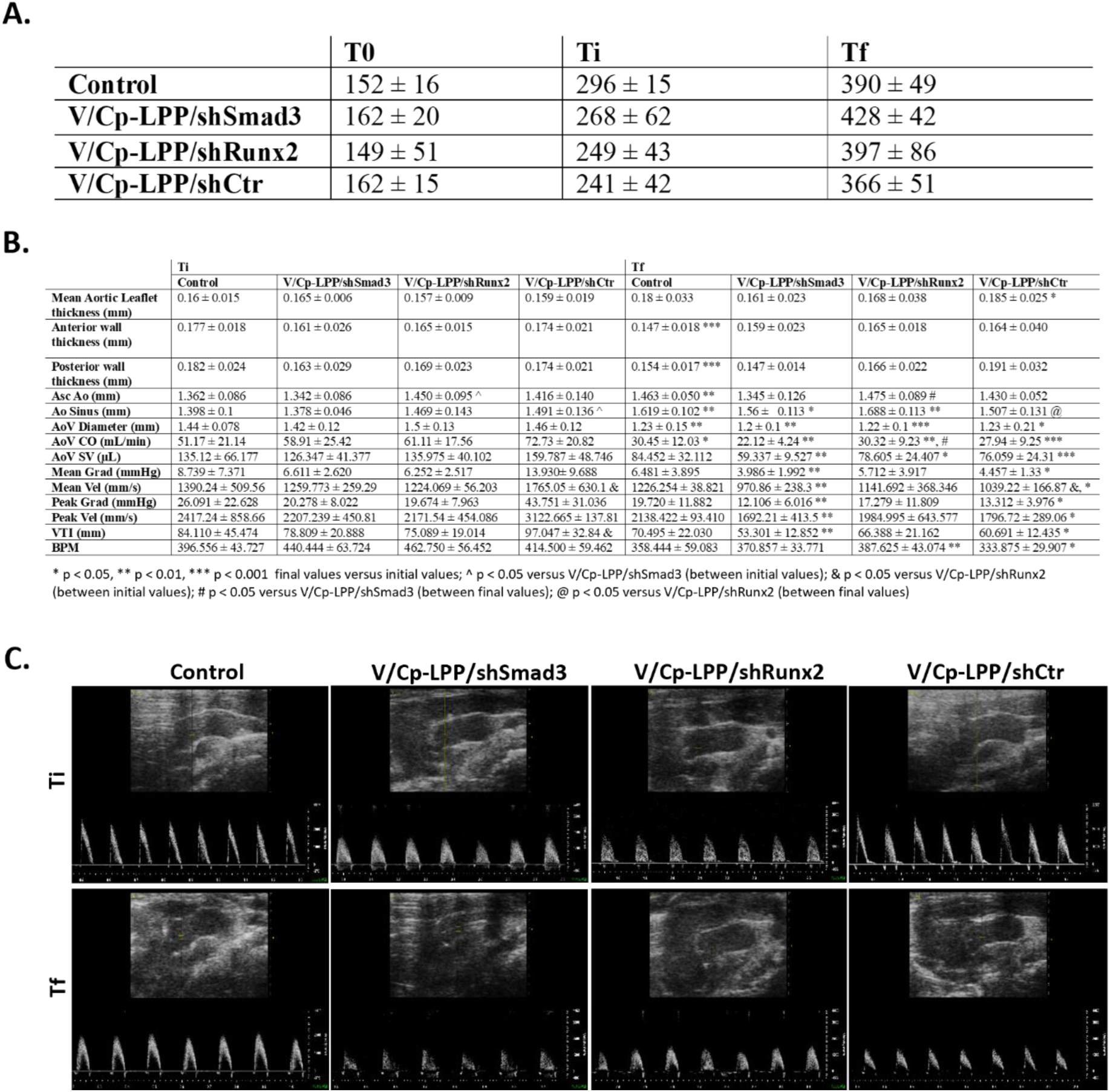
**A.** Glycaemia variation measured in diabetic/hyperlipidemic ApoE-deficient mice injected with V/Cp-LPP/shSmad3, V/Cp-LPP/shRunx2, V/Cp-LPP/shCtr lipopolyplexes, and control mice. The results are represented as a mean ± S.D. of n = 9 mice/experimental group. Abbreviations: T0: values before STZ injections, Ti: values after STZ administration, Tf: values before sacrifice. **B.** Echocardiographic parameters measured in the mouse model of CAVD injected with V/Cp-LPP/shSmad3, V/Cp-LPP/shRunx2, V/Cp-LPP/shCtr lipopolyplexes, and PBS, at two intervals after diabetes induction (initial, Ti) and before sacrifice (final, Tf). The data are expressed as mean ± S.D. (n = 9 mice/experimental group). Statistical significances (Sidak’s multiple comparisons test): * p < 0.05, ** p < 0.01, *** p < 0.001 final values versus initial values; ^ p < 0.05 versus V/Cp-LPP/shSmad3 (between initial values); & p < 0.05 versus V/Cp-LPP/shRunx2 (between initial values); # p < 0.05 versus V/Cp-LPP/shSmad3 (between final values); @ p < 0.05 versus V/Cp-LPP/shRunx2 (between final values). Abbreviations: Asc Ao: Ascending Aorta, Ao Sinus: Aortic Sinus, AoV Diameter: Aortic Valve Diameter, AoV CO: Aortic Valve Cardiac Output, AoV SV: Aortic Valve Stroke Volume, Mean and Peak Grad: Mean and Peak Gradient, Mean and Peak Vel: Mean Peak Velocity, VTI: Velocity Time Integral, BPM: Beats Per Minute. **B.** Representative PW-Doppler mode images of the aortas and aortic valves of diabetic/hyperlipidemic ApoE-deficient mice injected with V/Cp-LPP/shSmad3, V/Cp-LPP/shRunx2, V/Cp-LPP/shCtr lipopolyplexes, and PBS, taken after diabetes induction (initial, Ti) and before sacrifice (final, Tf).

Hemodynamic parameters were evaluated by echocardiography at two time points: Ti, the initial stage after diabetes induction, and Tf, the final stage before animal sacrifice (**Figure 5B**). Representative PW-Doppler mode echocardiographic images are shown in **Figure 5C**. Echocardiographic measurements of the aortic valve revealed leaflet thickness ranging from 1.6 to 1.85 mm across all the experimental groups at both Ti and Tf, exceeding the typical measurement of approximately 1.2 mm observed in C57BL/6 mice (**Table S3**). The thickness of both the anterior and posterior aortic walls remained unchanged across the experimental groups, with a mean value significantly higher than the measurements observed in the aorta of C57BL/6 mice (**Table S3**). The values for the Aortic Sinus (Ao Sinus) increased significantly, while the values of Aortic valve Diameter (AoV Diam) decreased significantly by the end of the investigation in both the control group (mice injected with PBS) and mice treated with lipopolyplexes. The functional parameters Cardiac Output (AoV CO) and Stroke Volume (AoV SV) significantly decreased by the end of the investigation period, regardless of the treatment groups (V/Cp-LPP/shSmad3, V/Cp-LPP/shRunx2 and V/Cp-LPP/shCtr). The Mean Gradient and Velocity, Peak Gradient and Velocity and Velocity Time Integral (VTI) significantly decreased at the end of the experiment in mice injected with V/Cp-LPP/shSmad3, by ∼ 40%, 23%, 40%, 23% and 32% (** p < 0.01), respectively. A significant decrease in these parameters was observed between the final and initial measurements after treatment with V/Cp-LPP/shCtr, with decreases of ∼ 68%, 41%, 70%, 42%, and 37% (*p < 0.05).

### V/Cp-LPP/shSmad3 and V/Cp-LPP/shRunx2 treatments suppress Smad3 and Runx2 mRNA expression in organs of the mouse model of CAVD

To investigate the systemic effects of lipopolyplexes in mice with CAVD, organ-specific mRNA levels of Smad3 and Runx2 were quantified following treatment. Administration of V/Cp-LPP/shSmad3 in the mouse model of CAVD led to a 40% reduction in mRNA-Smad3 level in the lungs and kidneys and a 50% decrease in the liver (**Figure S1B)**. In contrast, V/Cp-LPP/shRunx2 treatment reduced mRNA-Smad3 expression by approximately 20% in the lungs and 40% in the kidneys compared to control mice (PBS-injected), though these changes did not reach statistical significance. However, in the liver, mRNA-Smad3 levels declined significantly by 40% relative to controls. Regarding mRNA-Runx2 expression, treatment with V/Cp-LPP/shSmad3 resulted in a downregulation of 25% in the lungs, 30% in the kidneys, and 40% in the liver. Similarly, V/Cp-LPP/shRunx2 administration led to an approximately 25% reduction in all organs compared to controls. Treatment with V/Cp-LPP/shCtr lipopolyplexes had no impact on mRNA-Smad3 and mRNA-Runx2 levels in the lungs and kidneys; however, it promoted an increase in their expression in the liver.

### The effects of V/Cp-LPP/shSmad3 and V/Cp-LPP/shRunx2 treatments on the collagen deposition in the aortic valve of the mouse model of CAVD

We further evaluated the effect of V/Cp-LPP/shSmad3 and V/Cp-LPP/shRunx2 administration on the collagen deposition in the aortic valve of the mouse model of CAVD. Masson’s Trichrome staining revealed substantial collagen deposition in the aortic valves of all experimental groups (**Figure S2A**), especially at the aortic commissures (the site where two valve leaflets attach to the aortic wall), and a much lesser deposition in the leaflets. Mice treated with either lipopolyplexes did not influence the distribution of the collagen fibers; in the pulmonary, hepatic, and renal tissues, the Masson’s Trichrome staining did not reveal abnormalities regardless of the experimental group (**Figure S2B**). However, the stroma of the splenic tissue exhibited a considerable amount of blue-stained collagen fibers in all experimental groups. In addition, the perivascular areas were characterized by a large deposition of collagen fibers.

### Safety evaluation: organ histology and plasma biomarkers reflecting the liver and kidney function

#### Plasma parameters

After collection, the mice’s plasma samples were examined for lipid profile and hepatic and renal function markers. The treatment of diabetic/hyperlipidemic ApoE-deficient mice with VCAM-1/ Collagen IV double-targeted lipopolyplexes for two weeks did not affect the plasma hepatic and renal parameters. On the contrary, V/Cp-LPP/shRunx2 significantly lowered the concentration of circulating cholesterol and triglycerides (** p < 0.01), as compared to control mice injected with PBS (**Figure 6A)**. Fluorescence microscopy of Nile Red-stained hepatic tissue (green) was used to visualize the lipid deposition (**Figure S3A**). After quantification of the dye’s fluorescence, we determined that the mice injected with V/Cp-LPP/shRunx2 lipopolyplexes exhibited a slightly lower level of lipids (∼ 15%), but with no statistical significance (**Figure S3B**).

**Figure 6.**
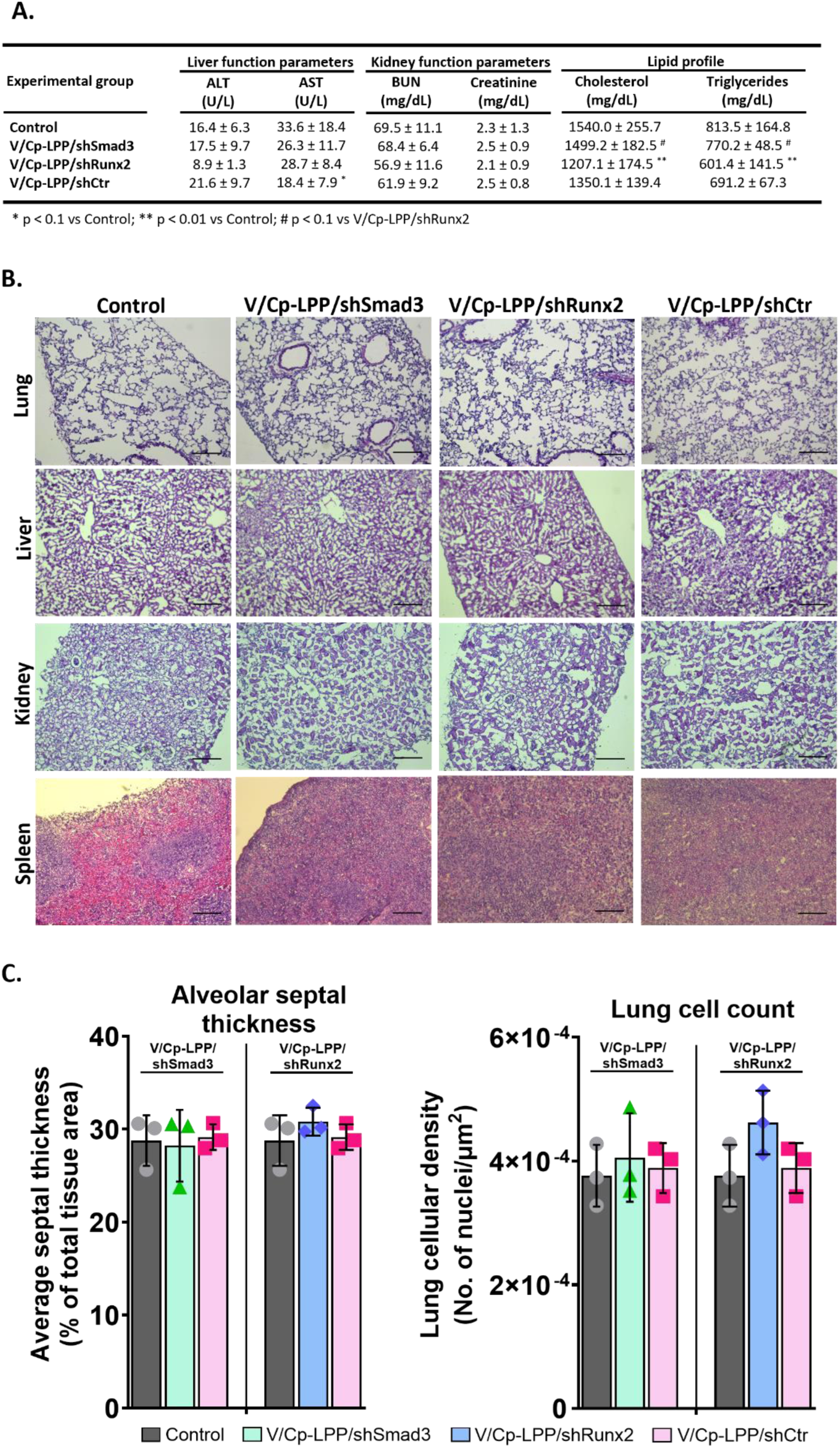
**A.** Levels of biochemical plasma parameters. Data are mean ± S.D. (n = 9 mice for each group). Statistical significance (one-way ANOVA with the Tukey post hoc test): * p < 0.1, ** p < 0.01 vs Control; # p < 0.1 vs V/Cp-LPP/shRunx2. **B.** Representative microscopy images of hematoxylin-eosin-stained organs harvested from diabetic/hyperlipidemic ApoE-deficient mice injected with V/Cp-LPP/shSmad3, V/Cp-LPP/shRunx2, V/Cp-LPP/shCtr lipopolyplexes or control mice. Scale bar: 200 µm. The image processing was performed using Fiji Is Just ImageJ (NIH) freeware program version 1.8.0. **C.** Alveolar wall thickness and lung cellularity measurement on images taken using a 20x objective (n = 3 mice for each group, ten random fields per mouse, illustrated as mean on the graphic were used for analysis using ImageJ 1.53c Macros described in Supplementary Material.

#### Histology

Tissue cryosections stained with hematoxylin-eosin were analyzed to assess histological changes in the lungs, liver, kidneys, and spleen of the mice treated with lipopolyplexes (**Figure 6B**). No obvious pathological lesions, such as edema, necrosis, and inflammatory infiltration, were detected in all the investigated experimental groups. No evidence of increased alveolar septal thickness or inflammatory cell infiltrates (**Figure 6C**) was observed in the examined lung sections. These data indicated that the lipopolyplexes did not induce structural or functional alterations in the liver, lungs, kidneys, and spleen.

## Discussion

The pathogenesis of CAVD is complex, involving many interconnected signaling networks that drive its progression.^29^ The activation and endothelial-to-mesenchymal transition (EndMT) of valvular endothelial cells (VEC) are critical events in the early stages of the disease, while valvular interstitial cell (VIC) osteodifferentiation and subsequent calcification drive disease progression. Many clinical trials aim to identify promising therapeutic strategies for CAVD. Still, these strategies have not yet translated into proven pharmacologic treatments for CAVD. The SALTIRE II RCT (NCT02132026) clinical trial focused on the effect of alendronate and denosumab in blocking the progression of valve stenosis.^30^ Based on the hypothesis that these drugs inhibit the receptor activator of nuclear factor κB ligand (RANKL), which plays a key role in VIC osteogenic differentiation and calcific nodules formation, their administration was expected to slow the progression of aortic stenosis.^31^ Despite 24 months of treatment, echocardiography and computed tomography (CT) scans revealed no significant improvement in cardiac function.^32^ The ARBAS RCT study (NCT04913870^33^) is a randomized controlled trial designed to slow the progression of valve calcification by targeting angiotensin receptors. Although angiotensin receptor blockade does not directly attenuate valve calcification, it alleviates fibrotic remodeling by reducing macrophage infiltration and downregulating IL-6 mRNA expression within the valve tissue.^34^ Studies conducted in eNOS−/− mice and a rabbit CAVD model have demonstrated that evogliptin, a dipeptidyl peptidase-4 (DPP-4) inhibitor, effectively decreases collagen I and fibronectin-1 expression within the aortic valve, thus supporting its potential therapeutic benefits in CAVD management. Given these data, evogliptin is currently evaluated for its effects in a phase 2 clinical study to assess its safety and efficacy in the treatment of CAVD (NCT05143177).^35^

We envisaged and evaluated a nanomedicine-based therapeutic strategy for CAVD, aiming to enhance targeted intervention and optimize disease management. We demonstrate here that our novel nanosystem, namely lipopolyplexes targeting both VCAM-1^15^ and collagen IV,^12^ accumulate in the aortic valve of a diabetic/ hyperlipidemic ApoE-deficient mouse model of CAVD. To enhance accumulation and retention within affected valve leaflets and facilitate uptake by VEC and VIC, we synthesized double-targeted lipopolyplexes functionalized with two peptides: one specific for VCAM-1 and the other for collagen IV. Physicochemical analysis of the dual-targeted lipopolyplexes encapsulating shRNA against Smad3 and Runx2 demonstrated a homogenous and stable colloidal solution, with a hydrodynamic diameter of ∼200 nm and a Zeta potential below ‒25 mV, parameters that remained unchanged over three weeks. Double-targeted lipopolyplexes, designed to carry a plasmid encoding a fluorescent protein, successfully facilitated efficient transfection of aortic valve leaflets in the mouse model of CAVD.

Then, we tested our hypothesis that repeated *i.v.* administration of lipopolyplexes carrying shRNA plasmids specific for Smad3 and Runx2 may exert a beneficial therapeutic effect on key molecules implicated in valve calcification, potentially influencing disease progression. We demonstrate that four administrations of either V/Cp-LPP/shSmad3 and V/Cp-LPP/shRunx2 effectively downregulate both gene and protein expression of Smad3 and Runx2 in the aortic valve leaflets (qRT-PCR and immunochemistry analysis). Moreover, we report a reduced level of Runx2 when Smad3 was downregulated using V/Cp-LPP/shSmad3 administration in CAVD mice as compared with untreated mice or mice treated with V/Cp-LPP/shCtr. It was previously reported that Smad3 transcription factor plays a crucial role in regulating Runx2 expression, as evidenced by data showing that heterozygous deletion of Smad3 in Klotho-deficient mice leads to the inhibition of Runx2, BMP2, and Alk2 (BMP Type I receptor) gene expression.^36^ These insights highlight the significant impact of Smad3 on osteogenic signaling pathways. We report here a reduced expression of osteogenic molecules such as ALP, OSP, and OSC in the aortic valve of animals treated with V/Cp-LPP/shSmad3. Moreover, we detected a reduction in ALP enzymatic activity. In contrast, treatment with V/Cp-LPP/shCtr lipopolyplexes did not lead to any significant changes in the levels of the investigated molecules. This suggests that an active mineralization process is ongoing in the valve tissue of untreated mice or control mice treated with dual-targeted lipoplexes carrying a control shRNA plasmid. Interestingly, we found a reduction of Smad3 expression when the mice were treated with V/Cp-LPP/shRunx2. Also, as in the case of V/Cp-LPP/shSmad3 administration, this treatment reduces the osteogenic molecules expression (i.e., ALP, OSP, OSC, and the mesenchymal marker αSMA). In prior experiments, we demonstrated that downregulation of Runx2 via lipopolyplexes targeted to VCAM-1 or collagen IV ^12,15^ in cultured osteoblast-committed VIC significantly reduces the expression of osteogenic molecules, including OSP, BSP, and BMP2. These findings suggest a direct regulatory role of Runx2 in the activation of downstream osteogenic molecules.

Previous studies have identified Smad2/3 as upstream regulators of Runx2, with their silencing resulting in a marked reduction in Runx2 protein expression and suppression of AngII-induced Runx2 activation in human aortic smooth muscle cells.^37^ Moreover, it has been demonstrated that Runx2-Smad interaction is required for the subnuclear recruitment of Smads at the active transcriptional sites^38^ and for increasing the binding affinity of Smad2/3 to DNA.^39^ The Smad interacting domain (SMID) of Runx2 is located in the C terminus of the Runx2 molecule, which overlaps with the nuclear matrix targeting signal (NMTS).^38^ A sequence of three residues, His-Thr-Tyr in the SMID domain, is involved in Runx2-Smad interaction and in enhancing the transcription of osteogenic genes.^40^ A mutation in this sequence affects the transcriptional activity of osteogenic genes (ALP, OSP, BSP, and OSC) by almost 75%.^41^ These findings are further confirmed and extended by our *in vivo* results showing that silencing Smad3 reduced the expression of Runx2 in the aortic valve, emphasizing the interaction between Smad3 and Runx2. While Runx2 expression was reported to be regulated by the Smad2/3 pathway, the direct impact of silencing Runx2 on Smad3 expression was not established before. In this study, we showed that when Runx2 is silenced using V/Cp-LPP/shRunx2, besides a reduced expression of Runx2, the Smad3 expression is also decreased, together with a reduction of osteogenic molecules expression. The co-dependent regulatory interplay between Runx2 and Smad3 was further confirmed and extended in our study using *in vitro* human osteodifferentiated VIC treated with lipopolyplexes carrying Runx2. Specifically, Runx2 downregulation led to a reduction in mRNA-Smad3 expression and vice versa. These results demonstrate their mutual regulatory dependence in integrating independent signal pathways to turn "on" specific genes involved in valve calcification.

While investigating the systemic action of lipopolyplexes, we detected a significant reduction in mRNA-Smad3 expression in the liver and lungs of mice treated with V/Cp-LPP/shSmad3 and in the liver of mice injected with V/Cp-LPP/shRunx2. The mRNA-Runx2 levels were reduced in both the kidneys and liver of mice treated with V/Cp-LPP/shSmad3, as well as in the liver of mice treated with V/Cp-LPP/shRunx2. Silencing of Smad3 and Runx2 in peripheral organs of CAVD mice demonstrated therapeutic potential in attenuating tissue fibrosis. Importantly, the absence of detectable histopathological or functional alterations following lipopolyplex delivery further reinforces the safety and tolerability of this therapeutic approach.

An elevated concentration of plasmatic alkaline phosphatase (ALP) has been identified as a predictor of mortality in both diabetes patients^42^ and individuals with CAVD.^43–45^ Three clinical studies (ASSERT, ASSURE, and SUSTAIN) reported the correlation between major adverse cardiovascular events (MACE) and higher ALP levels in patients. Reducing serum ALP activity through treatment with apabetalone, an epigenetic modulator that inhibits the interaction between BET (bromodomain and extraterminal domain) proteins and acetylated histones, for up to 26 weeks, has been shown to reduce cardiovascular risk in patients with MACE.^46^ Our findings demonstrate that, in addition to the reduction of ALP expression in the aortic valves of mice injected with lipopolyplexes carrying shRNA-Smad3 or shRNA-Runx2 plasmids, a decrease of plasma ALP levels was also obtained.

Conflicting findings regarding the association between cardiovascular disease severity and OSC level were reported. A positive correlation between the higher level of OSC and atherosclerosis,^47^ aortic calcification and coronary heart disease complicated with diabetes was reported.^48^ However, other studies concluded that in myocardial infarction,^49^ coronary artery disease,^50^ aortic atherosclerosis^51^ and calcification,^52^ the serum level of OSC is lower than in control subjects. In this study, the mRNA-OSC and mRNA-OSP expression was significantly reduced in the aortic valves, whereas plasma levels remained unchanged following treatment with lipopolyplexes.

Lipopolyplexes used in these experiments demonstrated high biocompatibility, as evidenced by the absence of renal and hepatic toxicity, as well as a lack of necrosis or inflammatory cell infiltration in the examined organs. Moreover, cholesterol and triglycerides decreased significantly after mice treatment with V/Cp-LPP/shRunx2. It has been proven that Runx2 controls sterol and steroid metabolism in osteoblasts by interacting with Runx2 elements in the Cyp11a1 promoter for the synthesis of cytochrome P450 that cleaves cholesterol to pregnenolone.^53^ Also, Runx2 modulates a gene network involved in fatty acid and cholesterol metabolism in macrophages^54^ and osteoblasts from calvaria.^55^ Considering these aspects, it is safe to assume that, in our experiments, downregulation of Runx2 in the hepatic tissue contributes to the reduction of plasma cholesterol and triglycerides.

To summarize, lipopolyplexes carrying shRNA-Smad3 and shRNA-Runx2 plasmids have multiple cardiovascular protective effects at the molecular level in the murine CAVD model. Therapeutic intervention resulted in decreased plasma levels of ALP, cholesterol, and triglycerides; partial modulation of the EndMT pathway, evidenced by reduced αSMA expression; and marked suppression of valvular calcification pathways through downregulation of key osteogenic transcription factors and markers such as Runx2, Smad3, OSP, ALP, and OSC. Yet, the echocardiographic measurements of the valve function indicated no statistically significant changes between the control group (mice injected with PBS) and the experimental groups injected with lipopolyplexes at the end of the investigation period. This can be explained by the short-term treatment, which induces modifications primarily at the molecular level, as demonstrated by qRT-PCR and immunohistochemistry. Targeting key transcriptional factors that regulate signaling pathways in CAVD through nanoparticle-based therapy represents a promising and innovative approach for disease management. This strategy holds promise for mitigating disease progression and advancing the therapeutic interventions in CAVD.

## Conclusions

The dual-targeted lipopolyplexes, engineered to recognize both VCAM-1 and collagen IV, effectively transfect valvular cells and induce robust expression of the plasmid-encoded protein within the aortic valve leaflets of a CAVD mouse model. Therapeutic delivery of V/Cp-LPP/shSmad3 and V/Cp-LPP/shRunx2 lipopolyplexes led to effective downregulation of Smad3 and Runx2, resulting in reduced levels of osteogenic markers, osteopontin, alkaline phosphatase, and osteocalcin, as well as diminished αSMA expression in the aortic valve of treated mice. Notably, our study reveals Runx2 as a previously unrecognized upstream regulator of Smad3 expression, establishing a novel Runx2–Smad3 axis with significant implications for valve pathobiology and therapeutic intervention. In addition, lipopolyplexes administration significantly lowered plasma levels of alkaline phosphatase, cholesterol, and triglycerides, supporting improved systemic metabolic balance while preserving liver and kidney function parameters.

Taken together, the data validate VCAM-1/ Collagen IV dual-targeted lipopolyplexes as an effective and selective treatment platform for CAVD, facilitating localized and sustained molecular reprogramming of affected valve tissue.

## Compliance with Ethics Requirements

All Institutional and National Guidelines for the care and use of animals were followed. The experiments were approved by the Ethics Committee of the Institute of Cellular Biology and Pathology “Nicolae Simionescu” and by the National Sanitary Veterinary and Food Safety Authority, authorization no. 448/ 02.04.2019 and were performed according to the Romanian Law no. 43/2014 (Official Monitor, Part I, nr. 326, pages 2–4), which transposes the EU directive 2010/63/EU on the protection of animals used for scientific purposes.

## Declaration of Competing Interest

The authors declare that they have no known competing financial interests or personal relationships that could have appeared to influence the work reported in this paper.

## Acknowledgments

The authors acknowledge Dr. Mariana Pinteala and Dr. Cristina Mariana Uritu for providing the nanoconjugate C60-PEI used for the polyplexes synthesis. This work was supported by the Competitiveness Operational Programme 2014–2020, Priority Axis1/Action 1.1.4/, Financing Contract no.115/September 13, 2016/MySMIS:104362, by the European Union – NextGenerationEU and the Romanian Government through the National Recovery and Resilience Plan, Component 9 - Investment 8, HeartCure project, Financing Contract no. 760063/23.05.2023, application no. CF93/15.11.2022 and by the Romanian Academy. Conceptualization: Calin M.; Methodology: Voicu G., Mocanu C.A., Safciuc F., Rebleanu D., Anghelache M., Turtoi M.; Investigation: Voicu G., Mocanu C.A., Safciuc F., Rebleanu D., Anghelache M., Turtoi M.; Formal Analysis: Voicu G., Mocanu C.A.; Validation: Voicu G.; Visualization: Voicu G., Mocanu C.A.; Writing – original draft, review, and editing: Voicu G., Simionescu M., Manduteanu I., Calin M.; Supervision: Simionescu M., Calin M.; Project administration: Manduteanu I.; Funding acquisition: Manduteanu I., Calin M.

## Supplemental Material

Supplemental Methods

Tables S1–S3

Figure S1-S3

Major Resources Table S4

---

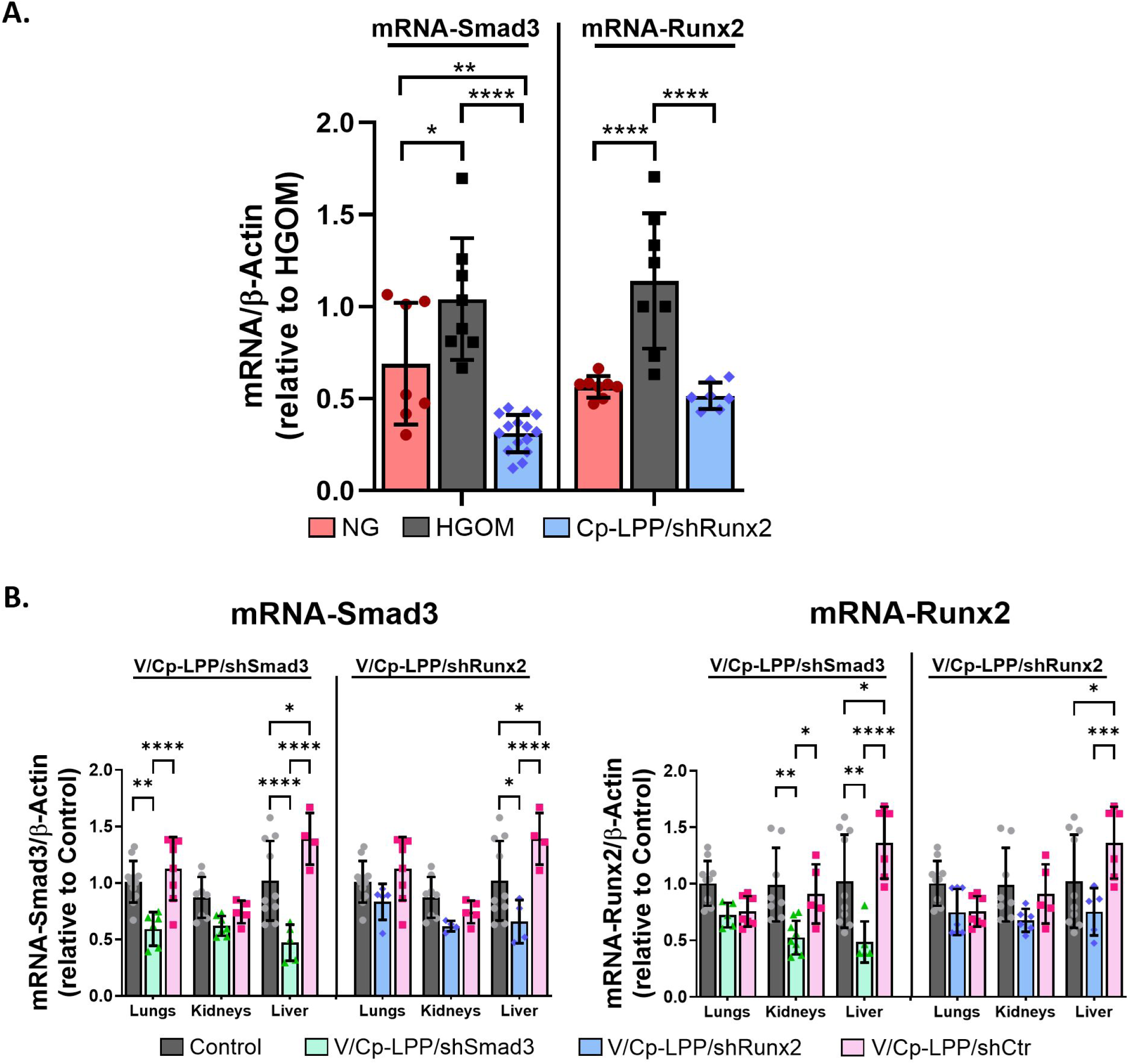

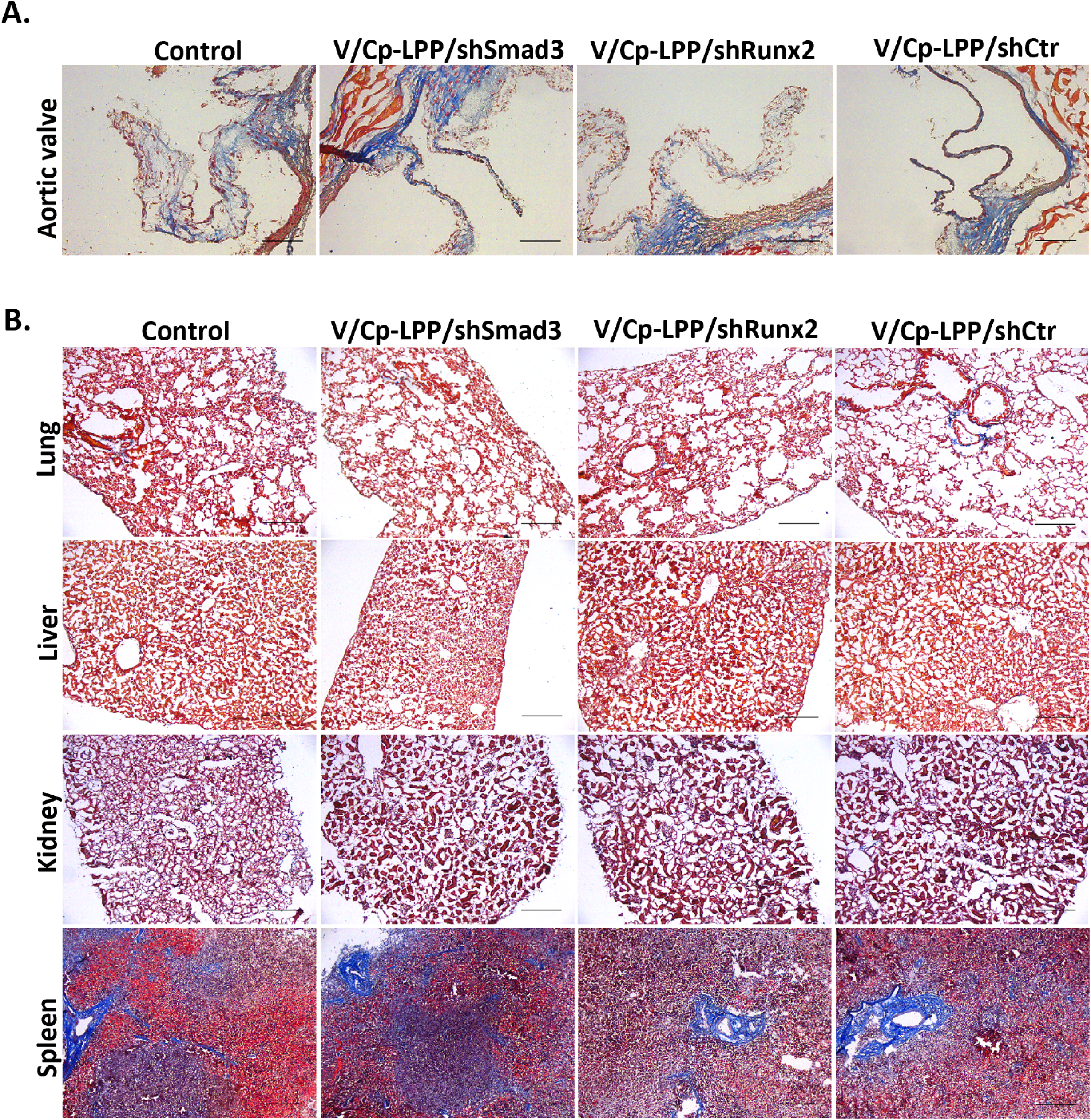

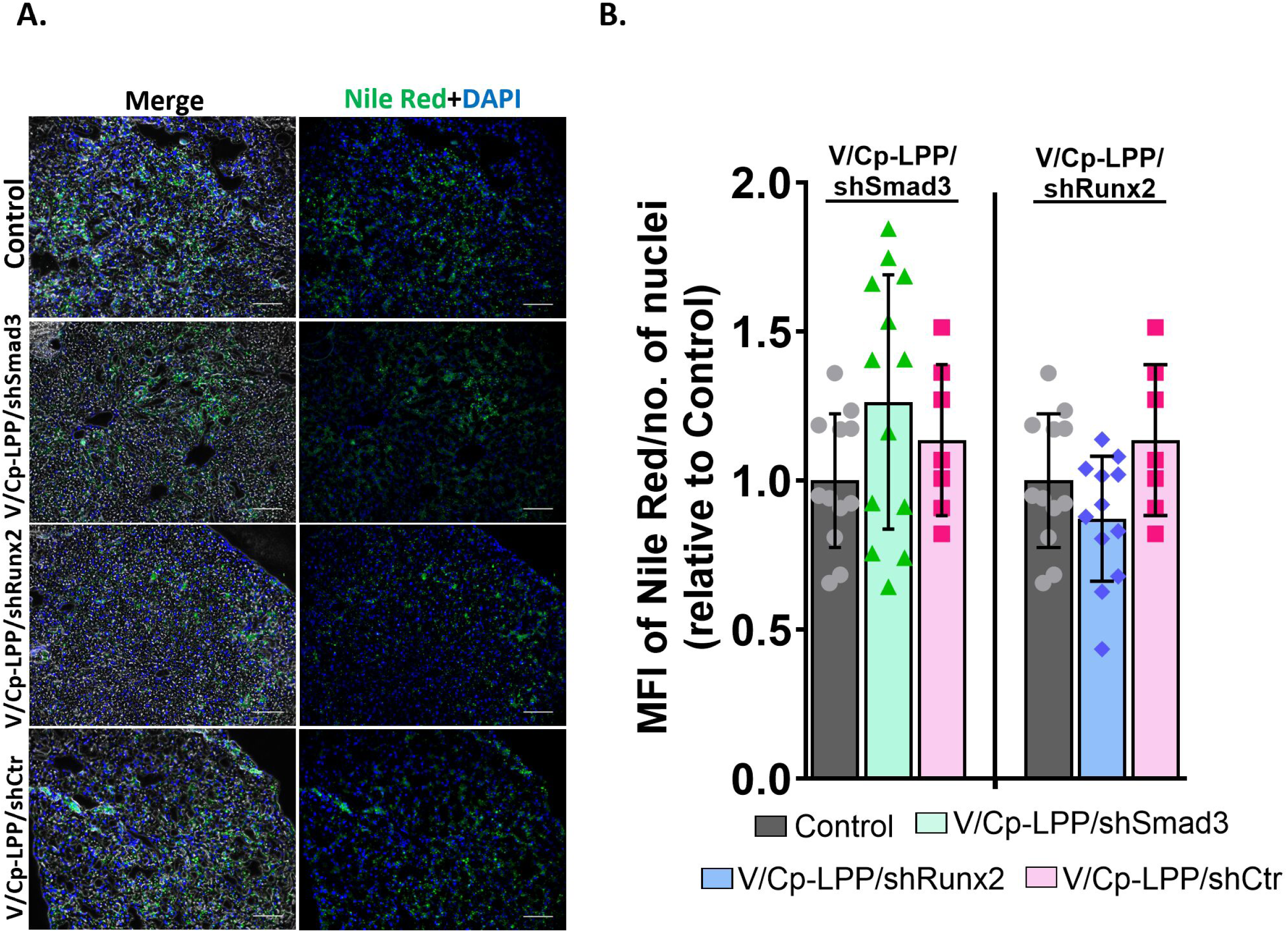

## Notes

### Competing Interest Statement

The authors have declared no competing interest.

